# Progressive reconfiguration of resting-state brain networks as psychosis develops: Preliminary results from the North American Prodrome Longitudinal Study (NAPLS) consortium

**DOI:** 10.1101/179242

**Authors:** Hengyi Cao, Yoonho Chung, Sarah C. McEwen, Carrie E. Bearden, Jean Addington, Bradley Goodyear, Kristin S. Cadenhead, Heline Mirzakhanian, Barbara A. Cornblatt, Doreen M. Olvet, Daniel H. Mathalon, Thomas H. McGlashan, Diana O. Perkins, Aysenil Belger, Larry J. Seidman, Heidi Thermenos, Ming T. Tsuang, Theo G.M. van Erp, Elaine F. Walker, Stephan Hamann, Alan Anticevic, Scott W. Woods, Tyrone D. Cannon

## Abstract

Mounting evidence has shown disrupted brain network architecture across the psychosis spectrum. However, whether these changes relate to the development of psychosis is unclear. Here, we used graph theoretical analysis to investigate longitudinal changes in resting-state brain networks in samples of 72 subjects at clinical high risk (including 8 cases who converted to full psychosis) and 48 healthy controls drawn from the North American Prodrome Longitudinal Study (NAPLS) consortium. We observed progressive reduction in global efficiency (*P* = 0.006) and increase in network diversity (*P* = 0.001) in converters compared with non-converters and controls. More refined analysis separating nodes into nine key brain networks demonstrated that these alterations were primarily driven by progressively diminished local efficiency in the default-mode network (*P* = 0.004) and progressively enhanced node diversity across all networks (*P* < 0.05). The change rates of network efficiency and network diversity were significantly correlated (*P* = 0.003), suggesting these changes may reflect shared underlying neural mechanisms. In addition, change rates of global efficiency and node diversity were significantly correlated with change rate of cortical thinning in the prefrontal cortex in converters (*P* < 0.03) and could be predicted by visuospatial memory scores at baseline (*P* < 0.04). These results provide preliminary evidence for longitudinal reconfiguration of resting-state brain networks during psychosis development and suggest that decreased network efficiency, reflecting an increase in path length between nodes, and increased network diversity, reflecting a decrease in the consistency of functional network organization, are implicated in the progression to full psychosis.

## Introduction

Substantial evidence has pointed to the disorganization of brain network architecture across the spectrum of psychiatric disorders involving psychotic symptoms. The most consistent findings in patients with psychotic disorders include altered network connectivity ^1-5^, network efficiency ^1, 5-11^ and network clustering ^1, 5, 6, 9, 10^, which disrupt information integration and segregation of brain systems, leading to abnormal topological structures of the brain. Moreover, altered network properties have been found to be related to genetic risk for psychotic disorders ^7, 9^, associated with severity of clinical symptoms ^5, 8^ and cognition ^1, 7, 11^, and predictive of antipsychotic response ^12, 13^. These lines of evidence suggest that changes in network integration and segregation may underlie the development of psychosis. However, whether these changes predict and potentially contribute to the onset of psychosis remains unclear.

Answering this question requires longitudinal observation of individuals at clinical high risk (CHR) prior to onset. Using this strategy, previous studies have identified progressive loss of gray matter in CHR subjects who converted to full psychosis compared with those who did not, involving regions that are critical to cognitive and social functioning, such as the dorsolateral and medial prefrontal cortex ^14-16^, temporal cortex ^16-18^ and cingulate cortex ^16, 17^. The progressive declines in gray matter volume and thickness are likely to be a result of excessive loss of neuropil (dendrites and synapses) during adolescence and early adulthood, which in turn, may lead to aberrant synaptic and neurotransmitter functioning that underlie abnormalities in brain connectivity and network configuration ^19^. Thus far, a critical piece of information is still missing in this model that directly links longitudinal changes in functional brain connectivity to the development of psychosis.

In the present study, using the data from the second phase of the North American Prodrome Longitudinal Study (NAPLS-2) consortium ^20^, we report on preliminary results of longitudinal changes in resting-state brain network architecture related to the conversion to psychosis. Here, 72 subjects at CHR, including 8 individuals who converted to psychosis, and 48 demographically comparable healthy participants underwent functional magnetic resonance imaging (fMRI) scans at both baseline and follow-up. We have previously shown that brain network measures derived from fMRI data are highly reliable both across time ^21^ and across scanner ^22^, making them particularly suitable for the multisite longitudinal design as used here. We hypothesized that converters would show progressive alterations and higher change rates in functional network properties compared to non-converters and controls, in particular measures assessing network connectivity, network integration and network segregation. Moreover, we predicted that these changes would be correlated with gray matter loss in converters.

## Methods and Materials

### Subjects

A sample of 120 subjects (72 clinical high risk (CHR) individuals, 48 healthy controls (HC, age 19.95±4.66 years, 28 males))) with available baseline and follow-up resting-state fMRI data was included in this study. During follow-up, 8 CHR subjects converted into psychosis (CHR-C, age 17.88±4.39 years, 5 males), and 64 subjects did not convert (CHR-NC, age 19.55±3.92 years, 39 males). The subjects were recruited as part of the NAPLS-2 consortium from eight study sites across the United States and Canada. The data included in the present study were drawn from a larger dataset with baseline scans as previously reported ^23^ (435 subjects in total, 27 converters, 245 non-converters and 163 controls). Notably, no significant differences were found between this longitudinally scanned subsample and the whole sample in terms of demographic and clinical measures (Table 1, *P* > 0.33). The study protocol was reviewed and approved by the institutional review boards at each site. All participants provided written informed consent.

**Table 1.** Characteristics of the studied sample

|  | **Converters**<br>**(n = 8)** | **Non-converters**<br>**(n = 64)** | **Controls**<br>**(n = 48)** | **P value** |
| --- | --- | --- | --- | --- |
| *Demographic variables* |  |  |  |  |
| **Age (years)** | 17.88±4.39 | 19.55±3.92 | 19.95±4.66 | 0.44 |
| **Sex (M/F)** | 5/3 | 39/25 | 28/20 | 0.91 |
| **Education (years)** | 10.88±2.85 | 12.15±2.55 | 13.02±3.24 | 0.09 |
| **IQ** | 101.25±15.71 | 110.27±16.29 | 114.53±14.20 | 0.08 |
| *Clinical variables* |  |  |  |  |
| **Baseline SOPS – positive** | 12.38±3.62 | 12.24±4.20 | 1.51±2.05 | <0.001 (overall)<br>0.92 (CHR-C vs CHR-NC) |
| **Follow-up SOPS – positive** | 18.00±6.74 | 7.32±4.90 | 0.81±1.71 | <0.001 (overall)<br><0.001 (CHR-C vs CHR-NC) |
| **Baseline SOPS – negative** | 9.50±5.12 | 12.24±6.37 | 1.28±1.85 | <0.001 (overall)<br>0.15 (CHR-C vs CHR-NC) |
| **Follow-up SOPS – negative** | 13.13±6.38 | 8.98±7.03 | 1.50±2.23 | <0.001 (overall)<br>0.15 (CHR-C vs CHR-NC) |
| **Baseline SOPS – disorganization** | 8.00±3.30 | 5.16±3.30 | 0.68±1.06 | <0.001 (overall)<br>0.01 (CHR-C vs CHR-NC) |
| **Follow-up SOPS – disorganization** | 9.75±4.80 | 3.98±3.37 | 0.56±1.07 | <0.001 (overall)<br><0.001 (CHR-C vs CHR-NC) |
| **Baseline SOPS – general** | 7.50±3.63 | 8.98±4.37 | 1.53±2.29 | <0.001 (overall)<br>0.28 (CHR-C vs CHR-NC) |
| **Follow-up SOPS – general** | 8.88±4.79 | 6.16±4.53 | 1.10±1.80 | <0.001 (overall)<br>0.16 (CHR-C vs CHR-NC) |
| **Baseline Antipsychotics (% medicated)** | 25 | 20 | 0 | <0.001 (overall)<br>0.40 (CHR-C vs CHR-NC) |
| **Follow-up Antipsychotics (% medicated)** | 50 | 17 | 0 | <0.001 (overall)<br><0.001 (CHR-C vs CHR-NC) |
| **Baseline Antipsychotics**<br>**(CPZ equivalent dosage)** | 70.31±131.26 | 44.89±109.25 | 0 | <0.001 (overall)<br>0.55 (CHR-C vs<br>CHR-NC) |
| **Follow-up Antipsychotics**<br>**(CPZ equivalent dosage)** | 50.00±111.80 | 44.76±120.96 | 0 | <0.001 (overall)<br>0.93 (CHR-C vs<br>CHR-NC) |
| *Cognitive variables* |  |  |  |  |
| **Baseline BVMT-R** | 19.00±9.05 | 27.01±5.80 | 27.66±4.99 | 0.001 (overall)<br>0.001 (CHR-C vs<br>CHR-NC) |
| **Follow-up BVMT-R** | 22.38±6.82 | 27.11±5.90 | 28.25±4.13 | 0.02 (overall)<br>0.06 (CHR-C vs<br>CHR-NC) |
| **Baseline HVLt-R** | 22.15±7.08 | 27.34±4.50 | 27.68±4.11 | 0.006 (overall)<br>0.008 (CHR-C vs<br>CHR-NC) |
| **Follow-up HVLt-R** | 23.63±4.63 | 27.49±4.73 | 28.77±3.74 | 0.008 (overall)<br>0.06 (CHR-C vs<br>CHR-NC) |
| *Image variables* |  |  |  |  |
| **Interscan interval**<br>**(months)** | 8.63±4.41 | 13.19±5.31 | 11.71±3.31 | 0.02 (overall)<br>0.03 (CHR-C vs<br>CHR-NC) |
| **Interscan FD Change** | 0.10±0.13 | 0.07±0.07 | 0.05±0.06 | 0.11 |

Abbreviations: SOPS = Scale of Prodromal Symptoms; CPZ = Chlorpromazine; BVMT-R = Brief Visuospatial Memory Test- revised; HVLt-R = Hopkins Verbal Learning Test- Revised; FD = Frame-wise Displacement. CPZ equivalent doses were calculated according to Leucht et al.<sup>44</sup>

The participants were evaluated using the Structured Clinical Interview for Diagnostic and Statistical Manual of Mental Disorders (SCID) ^24^ and the Structured Interview for Prodromal Syndromes (SIPS) ^25^ at each assessment point by clinicians. At each assessment point, prodromal symptom severity was quantified using the Scale of Prodromal Symptoms (SOPS) ^25^. Memory and learning abilities were evaluated using the Brief Visuospatial Memory Test-Revised (BVMT-R ^26^) and the Hopkins Verbal Learning Test-Revised (HVLT-R ^27^). The three groups were balanced with regard to demographic measures including age, sex, years of education and IQ (*P* > 0.08). The CHR-C and CHR-NC groups did not differ in the severity of positive and negative symptoms at baseline (*P* > 0.15), nor in antipsychotic dosages at both baseline and follow-up (*P* > 0.55). See Table 1 and Supplemental Materials for a detailed description of the sample.

### Imaging paradigm and data acquisition

All participants underwent a 5-min eyes-open resting-state scan. See Supplementary Materials for details on data acquisition.

### Data processing

The entire processing pipeline followed that of previously published work ^4, 21, 28^. In brief, mean time series for each of the 90 nodes defined by the Automated Anatomical Labelling (AAL) atlas ^29^ were extracted from the preprocessed data and further corrected for physiological and scanner noises. Pairwise Pearson correlation coefficients were calculated between the processed time series of each node, resulting in a 90 × 90 two-dimensional correlation matrix for each subject at each scan point (see Supplementary Materials for details).

We quantified two connectivity metrics describing the characteristics of the derived correlation matrices: node strength and node diversity ^1^. Node strength is the average connectivity between a given node and all other nodes in the network, reflecting how strongly the node is connected to others. Node diversity is the connectivity variance between a given node and all other nodes, reflecting how homogeneous the connectivity is for that node. These two metrics were then further averaged across the 90 nodes to generate a global measure.

To build weighted brain graphs, the derived correlation matrices were further thresholded into 31 densities ranging from 0.10 to 0.40 with an increment interval of 0.01. At each density, only the connections with correlation coefficients higher than the given threshold were kept as true internode connections in the matrix, thereby generating a weighted adjacency matrix. Three graph metrics were subsequently computed assessing the integration and segregation of the derived weighted networks: global efficiency, transitivity and small-worldness. Global efficiency is a measure of network integration, defined as the average inverse of the shortest path length between all pairs of nodes in the network. Transitivity quantifies network segregation as the normalized global measure of network clustering. Small-worldness is an index assessing the combination of network segregation and network integration. After computation, these measures were averaged across all densities to ensure that results were not biased by a single threshold.

As we have previously shown ^21, 22^, all examined connectivity and graph metrics are highly reliable across scanners and sessions, making the investigation of their longitudinal changes feasible. Here, following our previous work ^14^, we quantified the change rates for each of the examined measures for each subject. Change rate was defined as

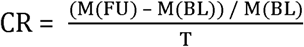

where M(FU) and M(BL) are network measures at follow-up and at baseline, respectively, and T is the time interval between the two scans (in month). As a result, change rate reflects the percentage of change for a given measure per month. This approach was preferred to repeated-measures analysis of variance (ANOVA) because the interscan interval varied across subjects.

### Statistical analysis

We employed an analysis of covariance (ANCOVA) model to test the differences in the examined metrics between the three groups at both baseline and follow-up. Here, the five baseline connectivity and graph metrics were entered as dependent variables in the baseline analysis, and their change rates were entered as dependent variables in the follow-up analysis. Group was given as independent variable, and age, sex, site, frame-wise displacement (for baseline analysis) and interscan change in frame-wise displacement (for follow-up analysis) were included as covariates. Significance was set at two-tailed *P* < 0.05 after false-discovery rate (FDR) correction for multiple comparisons of the five metrics.

All metrics with significant group differences were then examined for potential associations with structural, clinical and cognitive variables using Pearson correlations. Specifically, according to the psychosis onset model described previously, if progressive loss of gray matter drives changes in brain connectivity and brain networks, the change rates of cortical thickness and network measures should be highly correlated. We tested this hypothesis using data on cortical thickness for bilateral prefrontal cortex as previously described (see Cannon et al. ^14^ for details), whose change rates were calculated using the same formula as above. In addition to cortical thickness, we also probed potential associations of change rates of functional network measures with clinical symptoms and memory ability at baseline, as these baseline variables have previously been shown as predictive of conversion to psychosis ^30, 31^. Clinical symptoms were quantified by the summed scores of each domain in the SOPS (positive, negative, disorganization, general), visuospatial memory ability was assessed using the total recall score in the BVMT-R, and verbal memory ability was assessed using the total recall score in the HVLT-R.

## Results

### Group differences at baseline

At baseline, there were no significant differences between converters, non-converters and controls for all examined metrics (*P*_FDR_ > 0.10) in the present sample with both baseline and follow-up scans (120 subjects). The results remained non-significant when using the whole sample with baseline scans (435 subjects, *P*_FDR_ > 0.79).

### Group differences in change rates

We observed significant group differences in change rates of two examined metrics: global efficiency (*P*_FDR_ = 0.006) and average node diversity (*P*_FDR_ = 0.001). Post-hoc analyses showed that the change rate in global efficiency was significantly different between converters and controls (*P* = 0.004, Hedge’s *g* = 1.12), and approaching significance in the contrast between converters and non-converters (*P* = 0.052, Hedge’s *g* = 1.05). Here, unlike groups of controls and non-converters, both of which had positive mean change rates, as a group converters had a negative change rate (Figure 1A). Moreover, post-hoc analyses for average node diversity showed that converters had significantly larger positive change rates than both non-converters (*P* = 0.001, Hedge’s *g* =1.27) and controls (*P* < 0.001, Hedge’s *g* = 1.28, Figure 1B), suggesting that conversion to psychosis is associated with a progressive decrease in global efficiency but also a progressive increase in node diversity, reflecting less integration and consistency in functional network organization.

**Figure 1.**
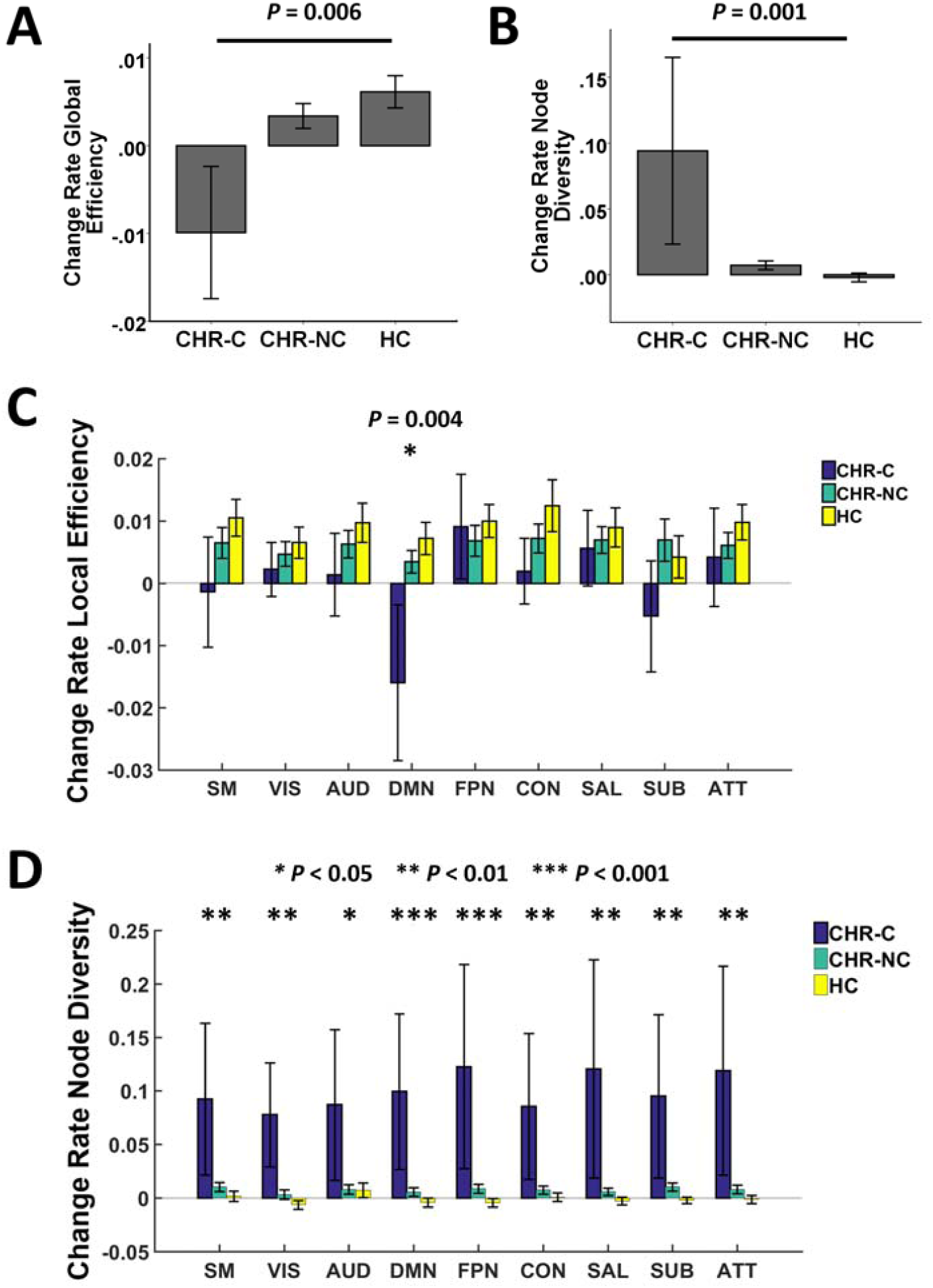
Association between change rates of network measures and conversion to psychosis. Converters showed progressively reduced global efficiency compared with non-converters and controls (Panel A), which was primarily driven by changes of local efficiency in the default-mode network (Panel C). In contrast, the progressive increase in node diversity observed in converters (Panel B) was distributed across the whole brain (Panel D). CHR-C = converters; CHR-NC = non-converters; HC = healthy controls; SM = sensorimotor network; VIS = visual network; AUD = auditory network; DMN = default-mode network; FPN = frontoparietal network; CON = cingulo-opercular network; SAL = salience network; SUB = subcortical network; ATT = attention network.

Given the results just summarized, we further investigated which brain systems were particularly involved in the differences between groups. For this, we calculated change rates of local efficiency and node diversity for each of nine well-established networks ^32^: sensorimotor, visual, auditory, default-mode, frontoparietal, cingulo-opercular, salience, subcortical and attention. Here, each of the 90 nodes were assigned to one or more systems based on Power et al. ^32^. Of note, Power’s study employed a different atlas with 264 nodes. As a result, some of the nodes in our study have been assigned to more than one network (in case different subregions of that node were allocated into different networks in Power’s study, see Table S1 for details). Change rates of local efficiency and node diversity were computed for each node and then averaged across nodes for each of the nine networks. The same ANCOVA model described above was used to test the group differences of the derived measures for each network, and significance was set at *P* < 0.05 after FDR correction for nine networks.

Our results revealed significant group differences in the change rate of local efficiency in the default-mode network (*P*_FDR_ = 0.004, Figure 1C). Similar to the pattern observed for global efficiency, on average converters showed a negative change rate in local efficiency in the default-mode network, compared with positive change rates among non-converters and controls. In contrast, no significant group differences were detected for other networks, suggesting that the progressive decrease of global efficiency was primarily driven by the local efficiency change in the default-mode network.

The analysis of change rate of node diversity demonstrated significant group differences for all of the nine examined networks (*P*_FDR_ < 0.046, Figure 1D). In particular, while non-converters and controls did not show significant changes in node diversity from baseline to follow-up (change rates approximately equaled to 0), converters showed significantly positive change rates for all nine networks, indicating that the progressive increase of node diversity in converters is not circumscribed but widely distributed across the whole brain.

### Association between change rates

We then investigated whether the change rates of these two measures reflected independent phenomena or shared neural mechanisms. Our analysis revealed significant negative correlations between change rate of network efficiency and change rate of network diversity both in the whole sample (*R* = -0.57, *P* < 0.001) and in the converter group (*R* = -0.89, *P* = 0.003, Figure 2), suggesting that these two functional alterations are highly dependent and possibly reflect shared underlying mechanisms.

**Figure 2.**
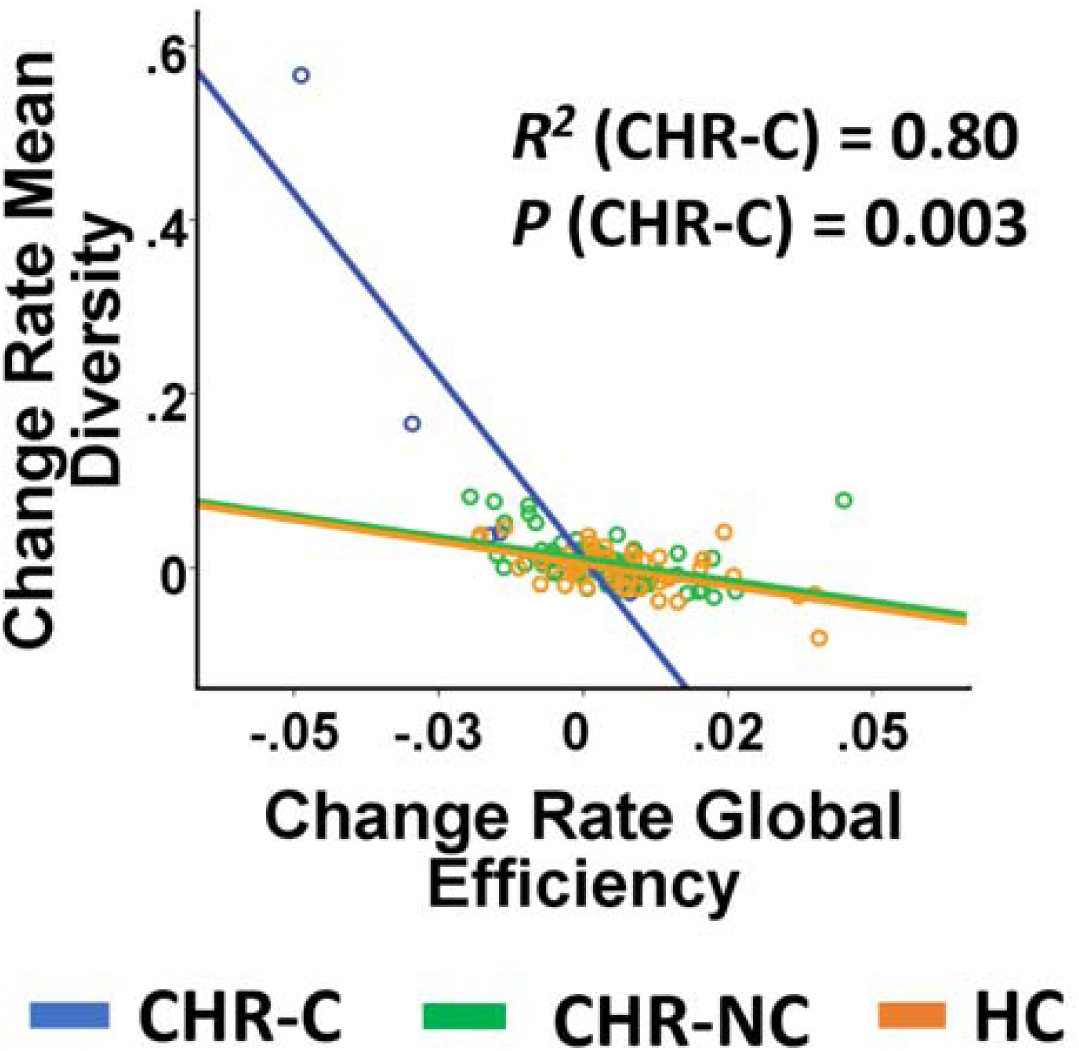
Association between change rate of network efficiency and change rate of network diversity stratified by outcome group. Two measures were significantly correlated with each other in converters and across all subjects.

### Association with cortical thickness

Correlation analyses identified significant negative correlations between change rates of cortical thickness in the prefrontal region and change rate of mean node diversity (*R* = -0.28, *P* = 0.002 and *R* = -0.40, *P* < 0.001 for left and right hemisphere, respectively, Figure 3A, 3B). Within the CHR-C group, change rates of both global efficiency and node diversity were significantly correlated with change rates of cortical thickness in both hemispheres (*R*^2^ > 0.56, *P* < 0.03 and *R*^2^ > 0.81, *P* < 0.003 for global efficiency and mean node diversity, respectively, Figure 3A-3D). In contrast, no significant correlations were found within the CHR-NC and HC groups.

**Figure 3.**
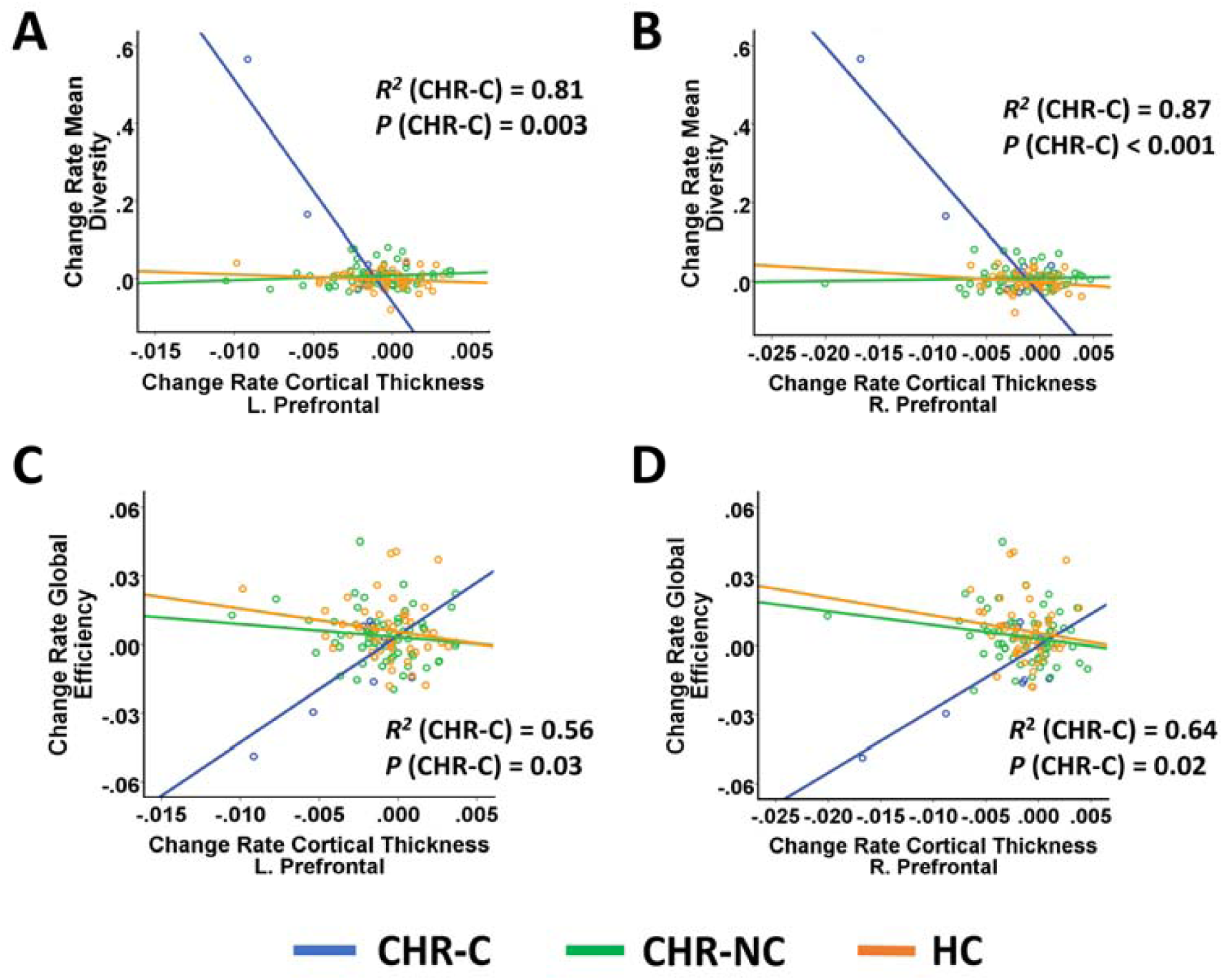
Association between change rates of network measures and change rates of cortical thickness in the prefrontal cortex stratified by outcome group. Change rates of both node diversity (Panel A & B) and global efficiency (Panel C & D) were significantly correlated with change rates of cortical thickness in bilateral prefrontal cortex in converters.

### Associations with clinical symptoms and memory ability

In the overall sample, a significant positive correlation was found between change rate of mean node diversity and baseline disorganization symptoms (*R* = 0.21, *P* = 0.02). Within the group of CHR-C, trend-level negative correlations were observed between change rates of both metrics and baseline negative symptoms, with relatively large effect sizes (all *R*^2^ > 0.23, Figure S1).

Change rates of both metrics were significantly correlated with BVMT-R total recall scores at baseline (*R* = 0.18, *P* = 0.04 for global efficiency and *R* = -0.37, *P* < 0.001 for mean node diversity, Figure 4A, 4B). The correlations were also significant when analyzed within the CHR-C group (*R*^2^ = 0.52, *P* = 0.04 for global efficiency and *R*^2^ = 0.64, *P* = 0.02 for mean node diversity) but not within the CHR-NC and HC groups. These findings suggest that baseline memory ability is predictive of the change rates of resting-state network measures in converters.

**Figure 4.**
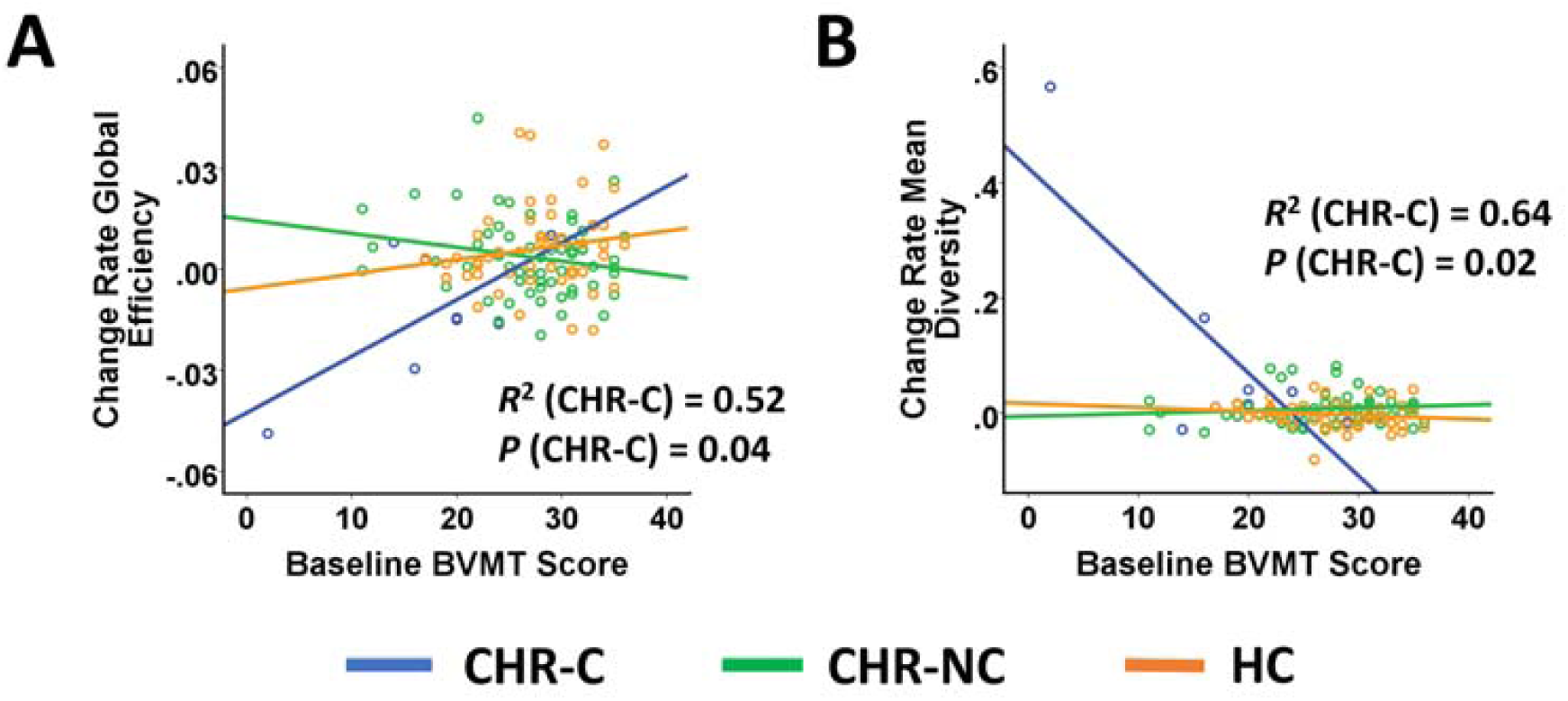
Association between change rates of network measures and baseline visuospatial memory ability as quantified by the Brief Visuospatial Memory Test (BVMT) total recall scores. Baseline BVMT scores were significantly correlated with change rates of global efficiency and node diversity in converters.

## Discussion

Using resting-state fMRI, this multisite longitudinal study found that CHR subjects who converted to psychosis showed a progressive decrease in global efficiency and increase in network diversity from baseline to follow-up at the point of conversion. These effects were primarily driven by progressive changes in local efficiency in the default-mode network (DMN) and changes in node diversity across the whole brain. Moreover, the identified alterations were highly correlated with each other and with progressive gray matter changes in the prefrontal cortex in converters, and could be predicted by subjects’ memory scores at baseline. These results provide preliminary evidence for functional network reorganization during the progression from a prodromal to fully psychotic state.

Deficits in network efficiency are among the most consistent findings in schizophrenia and other psychotic disorders and may serve as a transdiagnostic biomarker for the psychosis spectrum ^11^. Such deficiency has been reported in functional networks during resting state ^5, 6, 11^ and active tasks ^33, 34^, as well as in structural networks constructed from diffusion tensor imaging (DTI) ^7, 8^, and are associated with severity of psychotic symptoms ^5, 8^ and cognitive ability ^7, 11^, suggesting a robust network-based biological trait underlying psychosis. In line with these findings, our results further showed a progressive decrease of resting-state network efficiency in converters and strong correlations between this change and gray matter loss in the prefrontal cortex. These results suggest that increasingly diminished network integration is implicated in the development of psychosis, which may be explained, at least in part, by changes in gray matter structure. Moreover, several lines of evidence have further suggested that declines in network efficiency may relate to aberrant synaptic and neurotransmitter functioning. This interpretation is supported by the fact that the administration of a N-methyl-D-aspartic acid (NMDA) receptor antagonist can induce chronic disruption of brain global efficiency in animals that resembles findings in patients with schizophrenia ^35^, and that lower global efficiency at baseline is associated with worse response to antipsychotic medication at follow-up ^12^. Since glutamate receptor-mediated neural plasticity is pivotal to synaptic pruning ^36^, which can be further regulated by dopamine signaling ^37^, these findings echo the prevailing model interpreting the onset of psychosis and further suggest that altered network efficiency may participate in a cascade of events from excessive synaptic elimination to brain dysconnectivity that underlies psychosis development.

Intriguingly, the reduced efficiency is primarily driven by changes in the DMN, a brain system that is activated during rest but deactivated during attention-demanding tasks ^38^. A large body of work has shown that patients with psychosis have attenuated DMN deactivation during active tasks but enhanced within-DMN connection during resting state ^2, 39-41^, which may relate to exaggerated internally-focused thoughts and self-reference during rest and failure in suppression of these thoughts during task ^42^. These abnormalities have also been shown in subjects both at CHR ^43^ and at genetic high risk ^40^, suggesting a neurobiological trait that exists even before the onset of psychosis. Here, our results extend prior activation and connectivity findings by showing a critical association between deficits in DMN efficiency and the development of psychosis. It has been argued that the DMN dysfunction may be a consequence of diminished top–down regulation by the frontoparietal cognitive control network ^2, 42^, a neural mechanism that may as well lead to changes in DMN efficiency.

This study also found that conversion to psychosis is associated with a progressive increase in node diversity across all systems in the brain, which is negatively associated with change rate of cortical thickness in the prefrontal cortex and global efficiency. These findings suggest that the overall connectivity pattern becomes increasingly unstable and heterogeneous as psychosis develops, a phenomenon that may share the same underlying neural mechanisms with changes in cortical thickness and network efficiency. These results are in parallel with prior work revealing increased node diversity in patients with schizophrenia ^1^. Although only significant in the pooled sample, a positive correlation was found between change rate in node diversity and baseline disorganization symptoms. As a result, the increased heterogeneity in global connectivity may relate to disability of maintaining coherent and logical thoughts and difficulty in sustaining goal-directed attention in psychotic patients.

Change rates of both global efficiency and node diversity were significantly correlated with visuospatial memory scores at baseline in converters, suggesting that baseline cognitive functioning can predict progressive reconfigurations in resting-state brain networks in the ramp up to full psychosis. These findings are in line with previous work showing that lower network efficiency is associated with lower cognitive ability across different types of psychotic disorders ^11^. In addition, these findings are also consistent with our prior findings that baseline memory ability is a significant predictor of psychosis among CHR patients ^30, 31^. Together, these results suggest that resting-state network measures such as global efficiency and node diversity may act as potential mediators between impaired baseline cognitive functioning and the development of psychosis.

Several limitations of this study need to be clearly acknowledged. First, given the very small sample size of converters, the results reported in this paper must be considered preliminary, but merit further replication tests in larger cohorts. In particular, the observed large effect sizes in this sample were affected by a high leverage sample point, suggesting that the effect sizes for the detected group differences and correlations may be overestimated. However, the observation that individuals with the largest changes in network metrics had the largest changes in cortical thickness supports the validity of the data. Second, the follow-up scans for converters were acquired after the point of conversion. As a consequence, our study cannot be interpreted as isolating changes that occur prior to onset of psychosis. However, given the fact that progressive changes in network measures are associated with baseline cognitive ability, these changes are unlikely to be a secondary phenomenon. Third, medication effects cannot be ruled out from our sample. However, given that converters and non-converters did not show significant differences in antipsychotic dosages at both baseline and follow-up and the strong associations of changes in functional network properties with rate of prefrontal cortical thinning in converters, which has been shown to be independent of medication status in our previous work ^14^, the network findings here are unlikely to be solely caused by medication.

To sum up, individuals at CHR who converted to psychosis show progressive decreases in network efficiency and increases in network diversity, which were in turn associated with progressive loss of gray matter and predicted by baseline memory ability. These findings provide preliminary evidence for longitudinal reconfiguration of resting-state brain networks during the development of psychosis. Further work is encouraged to replicate these findings in larger samples and to investigate the predictive power of these findings for psychosis in independent cohorts.

## Acknowledgements

This work was supported by the National Institute of Health (NIH) grants U01 MH081902 to Dr. Cannon, P50 MH066286 to Dr. Bearden, U01 MH081857 to Dr. Cornblatt, U01 MH82022 to Dr. Woods, U01 MH066134 to Dr. Addington, U01 MH081944 to Dr. Cadenhead, R01 U01 MH066069 to Dr. Perkins, R01 MH076989 to Dr. Mathalon, U01 MH081928 to Dr. Seidman, and U01 MH081988 to Dr. Walker.

